# Comparative omics reveals unanticipated metabolic rearrangements in a high-oil mutant of plastid acetyl-CoA carboxylase

**DOI:** 10.1101/2023.08.31.555777

**Authors:** Amr Kataya, Jose Roberto S. Nascimento, Chunhui Xu, Matthew G. Garneau, Somnath Koley, Athen Kimberlin, Brian P. Mooney, Doug K. Allen, Philip D. Bates, Abraham J. Koo, Dong Xu, Jay J. Thelen

**Affiliations:** Department of Biochemistry, University of Missouri, Columbia, Missouri; Christopher S. Bond Life Sciences Center University of Missouri, Columbia, Missouri; Department of Electrical Engineering and Computer Science, University of Missouri, Columbia, Missouri; Institute of Biological Chemistry, Washington State University, Pullman, Washington; Donald Danforth Plant Science Center, St. Louis, Missouri; United States Department of Agriculture-Agricultural Research Service, St. Louis, Missouri

**Keywords:** fatty acid biosynthesis, acetyl-CoA carboxylase, seed oil, multi-omics

## Abstract

Heteromeric acetyl-CoA carboxylase (ACCase) catalyzes the ATP-dependent carboxylation of acetyl-CoA to produce malonyl-CoA, the committed step for *de novo* fatty acid synthesis. In plants, ACCase activity is controlled at multiple levels, including negative regulation by biotin attachment domain-containing (BADC) proteins, of which the *badc1/3* double mutant leads to increased seed triacylglycerol accumulation. Unexpectedly, the Arabidopsis *badc1/3* mutant also accumulates more protein. The metabolic consequences from both higher oil and protein was investigated in developing *badc1/3* seed using global transcriptomics, translatomics, proteomics, and metabolomics. Changes include: reduced plastid pyruvate dehydrogenase; increased acetyl-CoA synthetase; increased storage and lipid-droplet packaging proteins; increased lipases; and increased β-oxidation fatty acid catabolism. We present a model of how Arabidopsis adapted to deregulated ACCase, limiting total oil accumulation, and altering flux through pathways of carbon accumulation that presents possible targets for future bioengineering of valuable seed storage reserves.

## INTRODUCTION

Lipids are the most reduced form of carbon with importance for food, feedstock, and biofuel production, which has led to an ever-growing demand for improved oil crops (Thelen and Ohlrogge, 2002a). Anabolic and catabolic processes of metabolism are coordinated to result in the accumulation, and eventual turnover of oil in seeds. For instance, *de novo* fatty acid synthesis (FAS) in the plastid is coordinated with acyl flux through the cytosolic acyl-CoA pool, membrane lipid biosynthesis in the endoplasmic reticulum (ER), fatty acid modifications (desaturation and elongation), triacylglycerol (TAG) assembly in the ER, storage in oil bodies, and turnover with lipases, followed by β-oxidation of fatty acids in peroxisomes (reviewed in Li-Beisson et al., 2013). De novo FAS begins with the ATP-dependent carboxylation of acetyl-CoA to produce malonyl-CoA, catalyzed in plants by a plastidial acetyl-CoA carboxylase (ACCase) (Salie and Thelen, 2016). In most plants, there are two physically distinct ACCase forms: a plastid heteromeric ACCase (hetACCase) responsible for *de novo* FAS, and a homomeric ACCase localized in the cytoplasm, involved in the synthesis of very-long-chain fatty acids and flavonoids. HetACCase is a multisubunit complex comprised of four catalytic subunits: biotin carboxylase (BC), biotin carboxyl carrier protein (BCCP), and α- and β-carboxyltransferase (α- and β-CT) (reviewed in Salie and Thelen, 2016). Considering plastid fatty acid production can limit TAG accumulation (Baud and Lepiniec, 2010), increasing flux through FA synthesis has the potential to increase TAG content (Chapman and Ohlrogge, 2012).

Storage reserves within seeds depend on the supply of maternal precursors from vegetative parts of the plant and the developing embryo’s metabolic activities. Seeds mainly receive sucrose, glucose, and the amino acids glutamine, alanine, or asparagine as sources of carbon and nitrogen (Allen and Young, 2013; Alonso et al., 2007; Schwender and Ohlrogge, 2002). Manipulating expression levels of hetACCase subunits has resulted in diverse changes in fatty acid metabolism (Caroca et al., 2021; Chen et al., 2009; Thelen and Ohlrogge, 2002b; Wang et al., 2022). Complex mechanisms control *de novo* FAS, in part by regulating ACCase activity through light (Sasaki et al., 1997), downstream product inhibition by 18:1-acyl ACP (Andre et al., 2012), interactions with PII proteins (Bourrellier et al., 2010) that link nitrogen metabolism (Huergo et al., 2013), and regulatory subunits including the envelope-membrane localized carboxyltransferase interacting (CTI) (Ye et al., 2020) and the “biotin attachment domain-containing” (BADC) proteins (Salie and Thelen, 2016), the latter two regulatory components being three gene families in Arabidopsis. The BADC proteins are similar to the catalytic BCCP subunit but lack a biotinylation motif and the catalytically essential biotinyl Lys residue therein. The absence of biotin renders BADC catalytically inactive, and instead, BADC acts as apparent negative regulators for hetACCase by competing with BCCP for access to the ACCase holoenzyme, notably during the dark cycle (Salie and Thelen, 2016; Ye et al., 2020). In Arabidopsis, there are three BADC proteins, each of which have been demonstrated to act as hetACCase inhibitors, with knockout mutants for individual and combined *badc* genes resulting in increased seed oil content (Keereetaweep et al., 2018). Alternatively, BADCs have been hypothesized to facilitate the assembly hetACCase complex through associations of BC, BCCP α/b-CT stabilizing the larger quaternary-structural organization (Shivaiah et al., 2020), although in this model how the loss of BADCs lead to increased seed oil is speculative. Regardless, single and double knockout mutants of the BADC genes produces a higher seed oil phenotype, and highest oil is observed in the *badc1/3* knockout combination (Keereetaweep et al., 2018). Understanding the collateral, metabolic consequences of enhanced seed oil from mutations to plant-unique regulatory proteins for hetACCase has the potential to identify bottlenecks post FAS.

The current study details the metabolic consequences of enhancing seed oil content by hetACCase deregulation in the Arabidopsis *badc1/3* double knockout mutant. Using a combination of transcriptomics, proteomics, translatomics, and metabolomics alongside [^14^C]acetate pulse-chase metabolic tracing experiments in developing seed, the studies provide a comprehensive assessment of changes in seed storage metabolism. Alterations in metabolism in the *badc1/3* mutant accommodate and potentially counteract the enhanced flux to lipids provided by deregulation of ACCase.

## RESULTS

### Arabidopsis *badc1/3* seeds accumulate more oil and protein

To verify the effects of knocking out BADCs, storage reserve accumulation was analyzed in Arabidopsis *badc1/3* and compared to wild type (WT) seeds grown under the same conditions. The loss of both BADC1 and BADC3 resulted in a 17% increase in seed oil content (Fig. 1A), which is consistent with prior reports on BADC double mutants and confirms the beneficial effect of enhanced hetACCase activity (Keereetaweep et al., 2018; Ye et al., 2020). Further, the fatty acid profile for the *badc1/3* mutant demonstrated increases in both fatty acid elongation and desaturation (Supplementary Fig. 1). Interestingly, when seed protein was evaluated, the *badc1/3* mutant demonstrated a 10% increase in soluble proteins (Fig. 1B). To confirm increases in seed protein, extracted protein was analyzed by SDS-PAGE. The *badc1/3* seeds showed increases in both precursor and processed cruciferin (12S globulin) storage proteins (Fig. 1C).

**FIGURE 1:**
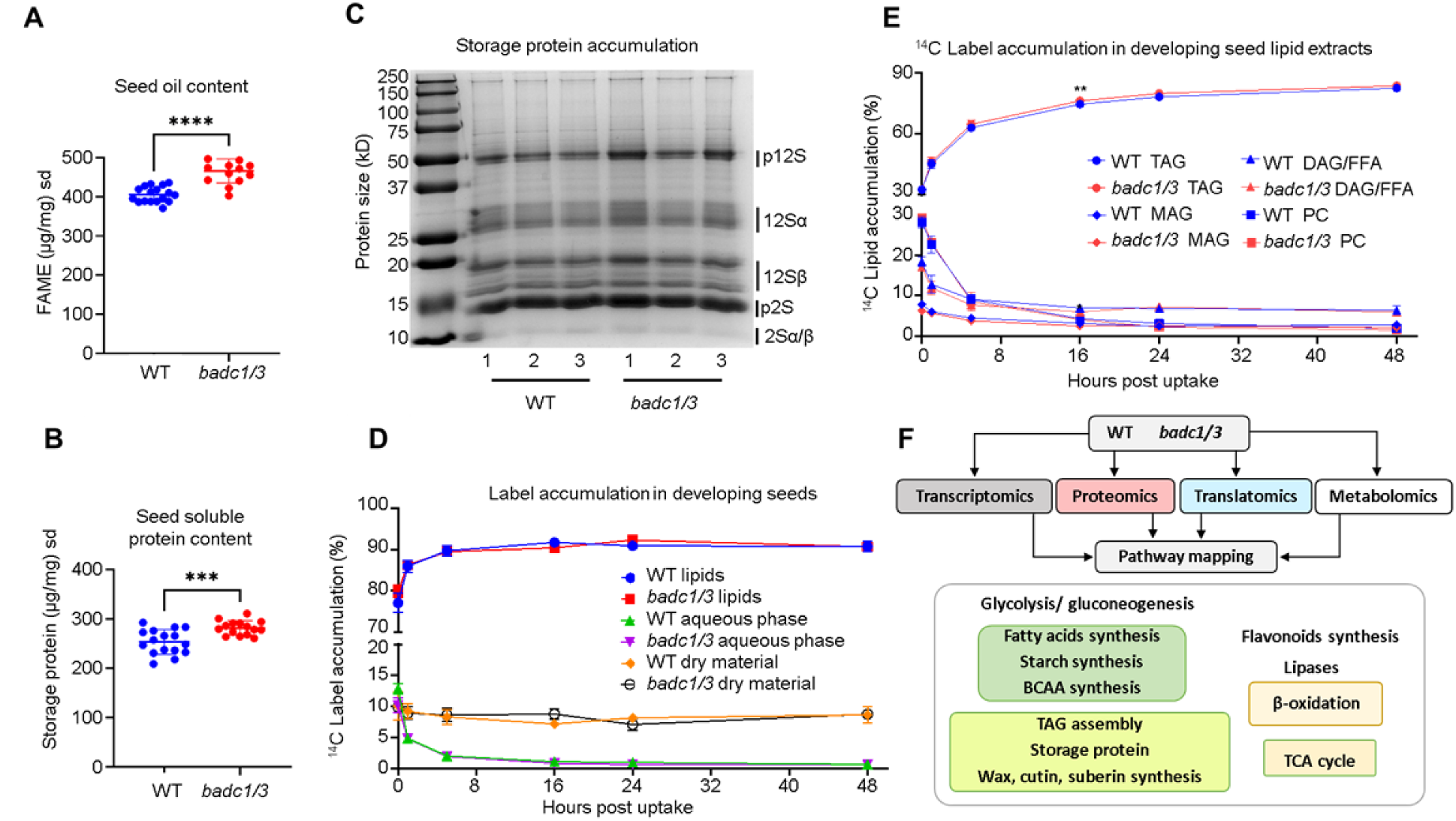
Seed oil and protein accumulation in mature *badc1/3* seeds, and [14C]acetate pulse-chase metabolic tracing in developing seeds at 9-10 days after flowering. Dry seeds from wild-type (WT) and *badc1/badc3* (*badc1/3*) Arabidopsis were analyzed for changes in oil (a) and soluble protein (b) (n≥16). Accumulation of storage proteins is compared from 3 representative WT and *badc1/3* samples on a Coomasie-stained 12% SDS-PAGE gel (c). Major storage proteins, including preprocessed 12S globulin/cruciferin (p12S) and 2S albumin (p2S) as well as processed α/β globulin (12Sα, 12Sβ) and albumin storage proteins (2Sα/β) labeled at right. Seed oil and protein values each represent a separate individual plant with included average and error bars which represent mean and standard deviation, respectively. The incorporation of [^14^C]acetate in developing embryos of WT and *badc1/3* knock-out line was used to determine the rate of fatty acid synthesis over 1 hour of pulse labeling, and the incorporation in different fractions followed during a 48 h chase. Label accumulation is shown in (d) the extracted lipid, aqueous, and remaining dry mass; as well as (e) the four most predominant lipid classes in developing seeds over the then entire chase period (n=4). Data are presented as a percentage of total radioactivity normalized to developing seed mass, data points represent average ± standard deviation. Asterisks represent significant differences from WT using a T-test (** P ≤ 0.01,*** P ≤ 0.001, **** P≤ 0.0001). (f) Summarized workflow used in multi-omics analysis.

### Isotopic tracing indicates increased seed oil in *badc1/3* is caused by relatively small increases in FAS and acyl partitioning to TAG during seed development

To initially confirm that the loss of BADC1 and BADC3, affected the rate of fatty acid synthesis (FAS) developing *badc1/3* seedlings were pulsed with [^14^C]acetate for 20 min. A comparison with WT detected an increase in ^14^C accumulation in the lipid fraction in the *badc1/3* seedlings consistent with increased incorporation of ^14^C into FAs in *badc1/3* seedlings (Supplementary Fig. S2). To elucidate the effect of the *badc1/3* mutation on FAS and lipid accumulation in developing seeds during the TAG accumulation stage, developing seeds were collected and the rate of fatty acid synthesis and acyl partitioning into membrane lipids and TAG were measured through a [^14^C]acetate pulse/chase experiment. Developing seeds at 9-10 days after flowering (DAF) were pulsed for 1 h with [^14^C]acetate, washed, and further incubated without the radiolabel during a 48 h chase (Fig. 1 D, E). At the end of the pulse (t = 0), the *badc1/3* labeling of the lipid fraction showed a higher trend than WT though it was not statistically significant (Fig. 1D). The analysis of ^14^C-fatty acid partitioning into different lipids (Fig. 1E) indicated that initial flux of newly synthesized fatty acids into PC (t = 0) was, on average, elevated. This also corresponded, on average, to more incorporation into TAG over the 48 h chase; however, most changes in lipid labeling were small.

### Integrating global transcriptomic, translatomics, and proteomics for a comprehensive assessment of *badc1/3* seed metabolism

The unexpected increase in storage protein, alongside enhanced oil accumulation, prompted multi-omics analysis to investigate the metabolic alterations resulting from knocking out BADC1 and BADC3 (Fig. 1F). Multi-omics analysis was performed on WT (Col-0) and *badc1/3* Arabidopsis developing seeds at 9-10 DAF, which represents the stage when developing Arabidopsis seeds rapidly accumulate storage reserves (Ruuska et al., 2002). Transcriptome deep-sequencing (RNA-Seq) resulted in nearly 40 million reads for each sample. The sequenced reads were mapped against Arabidopsis TAIR10 annotated genome, resulting in no less than 98.7% of total reads mapped to a total of 23,150 Arabidopsis genes. Among them, 2,802 genes were differentially expressed (q value < 0.05) (Supplementary Table 1). Translatome analysis consisted of the parallel deep-sequencing of mRNA engaged with ribosomes (TRAP-Seq), which was compared with the total mRNA deep-sequencing to calculate translation efficiency (TE), as previously described (Bailey-Serres et al., 2009; Kimberlin et al., 2021; Mazzoni-Putman and Stepanova, 2018; Reynoso et al., 2015; Xu et al., 2017). TE was determined for a total of 13,272 genes identified (Supplementary Table 2). A total of 668 genes were found to have TE changes (Z-score >2 or <-2) in *badc1/3* compared to WT. Proteomic analyses employed gel-free tandem mass tag (TMT) labeling with high coverage PASEF-LCMS on a timsTOF PRO. MS/MS spectra search against the Arabidopsis TAIR11 database identified a total of 4,033 proteins following the criteria of peptide spectrum-matches FDR 0.1% and ≥ 1 unique peptide per protein, with 513 differentially expressed proteins (p-value < 0.05) (Supplementary Table 3).

Parallel omics facilitated the comparative analysis of thousands of genes/proteins, providing a comprehensive assessment of the changes caused by metabolic perturbation. Correlating and interpreting data from parallel omics approaches is challenging because gene expression changes are frequently discordant with the proteome due to several factors, such as post-transcriptional and post-translational regulation (Fernie and Stitt, 2012; Hajduch et al., 2010). In the current study, changes in transcript and protein levels agreed for approximately 48% of cognate transcript/protein pairs, similar to a transcriptome/proteome profile of developing Arabidopsis seed (Hajduch et al., 2010). Utilizing the omics datasets, a comprehensive assessment of metabolic alterations was obtained (Fig. 2-5). Subcellular localization was mapped according to the SUBA5, Aralip, and TAIR databases (Supplementary table 4) (Hooper et al., 2017; Li-Beisson et al., 2013). Mapping was focused on differentially expressed genes and proteins (p/q-value < 0.05, log2foldchange ≥ 1, ≤ −1 for protein and transcript data, and Z-score ≥ 2, ≤ −2 for Trap-Seq data).

**FIGURE 2.**
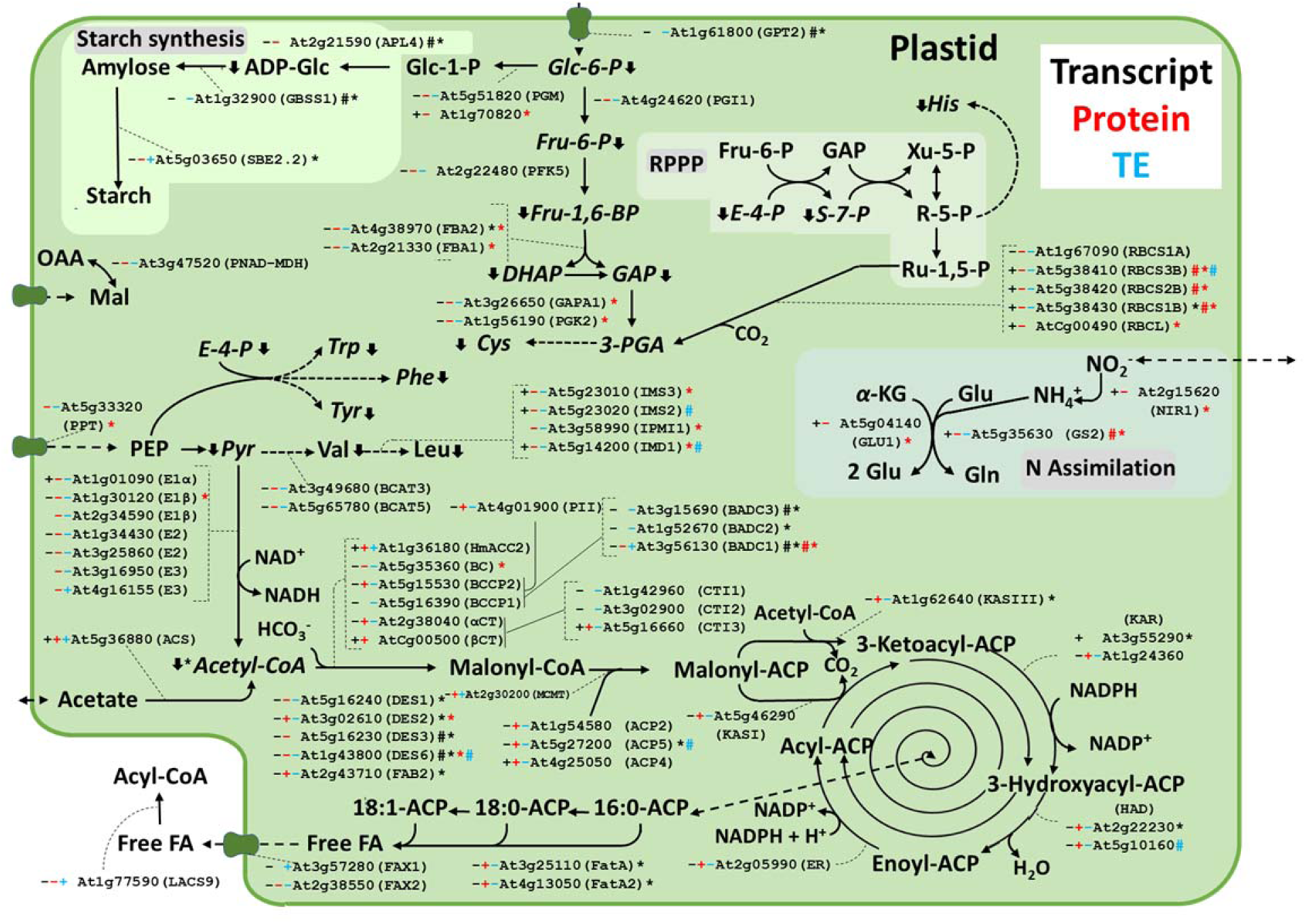
Global protein, transcript, and translation efficiency changes in plastids due to *badc 1/3* knockout in developing Arabidopsis seeds. The plus and minus signs represent upregulation and downregulation respectively in the *badc1/3* line. The black, red, and blue color represent transcript, protein, and translation efficiency data respectively. The number sign represents Log2FC ≥1 or ≤ −1 for transcript and protein, and the asterisk represents genes with q value ≤ 0.05 for transcript and p values ≤ 0.05 for protein, these are also color-coded. The blue number sign represents translation efficiency Z-score ≥ 2 or ≤ −2. The downward and upward arrows represent decreased and increased metabolite levels respectively. The asterisk next to the arrow represents p values ≤ 0.05 for the difference in metabolite levels between *badc1/3* mutant and WT. Abbreviations: glucose-6-phosphate/phosphate transporter 2 (GPT2); phosphoglucose isomerase 1(PGI1); ADP-glucose pyrophosphorylase large subunit (APL4); Granule bound starch synthase 1 (GBSS1); starch branching enzyme 2.2 (SBE2.2); phosphofructokinase 5 (PFK5); glyceraldehyde-3-phosphate dehydrogenase a subunit 1 (GAPA1); phosphoglycerate kinase 2 (PGK2); rubisco small subunit (RBCS); rubisco large subunit (RBCL); plastidic NAD-dependent malate dehydrogenase (PNAD-MDH); triose-phosphate∕phosphate translocator (TPT); phosphate/phosphoenolpyruvate translocator (PPT); pyruvate dehydrogenase E1 alpha subunit (E1α); pyruvate dehydrogenase E1 beta subunit (E1β); dihydrolipoamide acetyltransferase (E2); dihydrolipoamide dehydrogenase (E3); acetyl-CoA carboxylase 2 (HmACC2); biotin carboxylase (BC); biotin carboxyl carrier protein (BCCP); carboxyltransferase (α- and βCT); acetyl-CoA synthetase (ACS); biotin-attachment domain containing (BADC); ketoacyl-ACP Synthase I/II/III (KASI/II/III); ketoacyl-ACP reductase (KAR); hydroxyacyl-ACP dehydrase (HAD); enoyl-ACP reductase (ER); acyl carrier protein (ACP); stearoyl-ACP desaturase (DES); ABC acyl transporter (FAX1); acyl-ACP thioesterase (FaTA); long-chain acyl-CoA synthetase (LACS); acyl-CoA binding protein (ACBP).

### Omics data supports repression of plastid pyruvate dehydrogenase complex and induction of acetyl-CoA carboxylase in *badc1/3* developing seed

Expression of plastid-localized proteins (Fig. 2) pointed to increased FAS in the *badc1/3* mutant. BADC1 (At3g56130) was strongly reduced at both protein and transcript levels, being the most downregulated protein in both datasets, while BADC3 (At3g15690) was significantly reduced at the transcript level and not detected at the protein level, confirming BADC1 and 3 were both knocked out in the mutant. Most of the catalytic subunits of hetACCase were increased at the protein level (BCCP2, α-CT, β-CT), except for BC. Additionally, plastid-localized homomeric ACCase (HmACC2, At1g36180) was increased at the transcript, translation efficiency (TE), and protein levels. Enzymes downstream of ACCase, including malonyl-CoA-ACP malonyltransferase (MCMT, At2g30200), and most FAS complex proteins including ACPs 2, 4, and 5 (At1g54580, At5g27200, and At4g25050) were alternately changed at transcript and protein levels, but only slightly upregulated at the protein level. As acyl-ACPs are essential intermediates of FAS, the higher level of the total ACP pool could indicate enhanced FAS capacity (Huang et al., 2017). Thus, despite reduced transcript and TE for most FAS complex enzymes, the higher protein abundances were consistent with increased FA synthesis in *badc1/3*.

Each subunit to the plastid pyruvate dehydrogenase complex (pPDC) was downregulated at transcript, protein, and TE levels in the mutant (Fig. 2), while changes in acetyl-CoA synthetase (ACS, At5g36880) were increased in all three datasets, but the increase was marginal. Additionally, two aldehyde dehydrogenases involved in acetaldehyde conversion to acetate (Kirch et al., 2004; Wei et al., 2009), one cytosolic aldehyde dehydrogenase (ALDH2C4, At3g24503), and one mitochondrial (ALDH2B7, At1g23800), were upregulated at the transcript, protein, and TE levels (Fig. 5), indicating a possible alternative supply of carbon for FAS. The data suggested a putative shift from pyruvate to acetate as part of the carbon source for FAS in the mutant. Omics data on acetaldehyde synthesis was inconclusive (Fig. 5). Although pyruvate decarboxylase 1 (PD1) was downregulated at transcript level in *badc1/3*, PD2 (At5g54960) was not differentially expressed, and PD4 (At5g01320) was upregulated at transcript and protein level, though not statistically significant. The role of each PD gene in pyruvate conversion into acetaldehyde has been analyzed under low oxygen stress conditions (Kursteiner et al., 2003; Mithran et al., 2014), but not with increased oil mutants. Therefore, as our multi-omics did not collect conclusive data for most PD genes, we speculate pyruvate conversion into acetaldehyde is heightened, which would be consistent with the proposed putative shift from pyruvate to acetate as part of the carbon source for FAS in the increased oil content in the mutant.

Enzymes for plastid carbohydrate metabolism was mostly decreased (Fig. 2). The plastid glycolytic enzymes, including phosphoglucose isomerase 1 (PGI1, At4g24620), phosphofructokinase 5 (PFK5, At2g22480) fructose-bisphosphate aldolase (FBA, At4g38970, At2g21330, and At4g26530), glyceraldehyde 3-phosphate dehydrogenase (GAPA, At3g26650 and At1g12900), and phosphoglycerate kinase (At1g56190 and At3g12780) were reduced at the transcript, protein, and TE levels. The protein and transcript levels of RuBisCO were lower for both large (AtCg00490) and small subunits (At5g38410, At5g38420, At5g38430, and At1g67090). Transport of glycolytic intermediates into the plastid was also reduced, as indicated by the low expression levels of glucose 6-phosphate/phosphate transporter 2 (GPT2, At1g61800), and the phosphate/phosphoenolpyruvate (PEP) translocator (At5g33320). Plastid carbohydrate metabolism reduced at transcript level suggested transcriptional regulation of plastid glycolysis that was consistent with the reduction in transcription factor WRINKLED1 (WRI1, At3g54320) in the mutant (Table 1), which is a key regulator of most key genes involved in converting sucrose into FA (Kong & Ma, 2018). Downregulation of plastid glycolysis was consistent with the proposed shift from pyruvate to acetate as part of the carbon source for FAS in *badc1/3*. Furthermore, reduction in transcripts of plastid starch synthesis genes, including ADP-glucose pyrophosphorylase (APL4, At2g21590), granule bound starch synthase 1 (GBSS1, At1g32900), and starch branching enzyme (SBE2.2, At5g03650), suggested reduced plastidic carbohydrate metabolism and a change in carbon allocation from starch to oil.

**Table 1.**
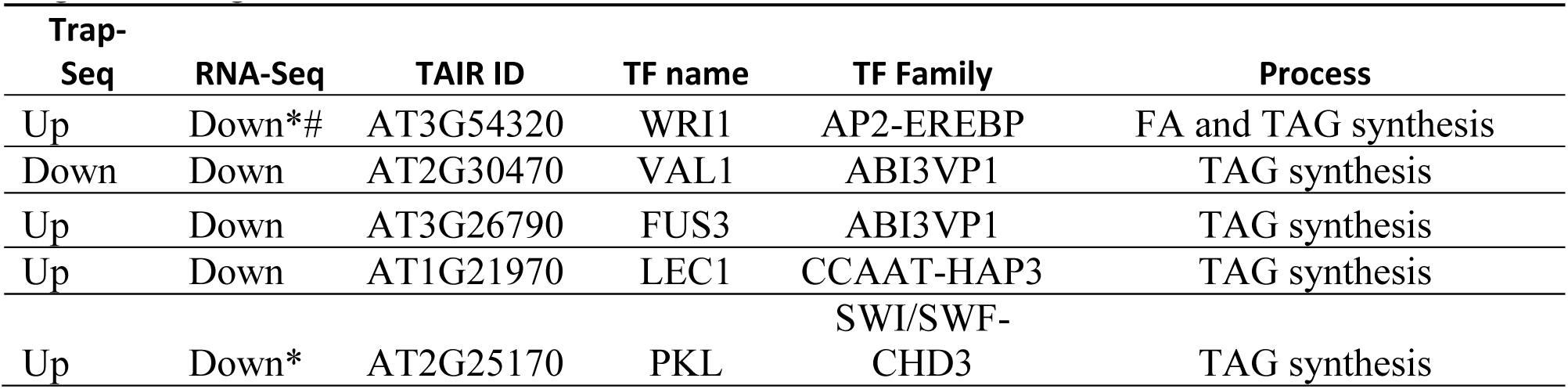
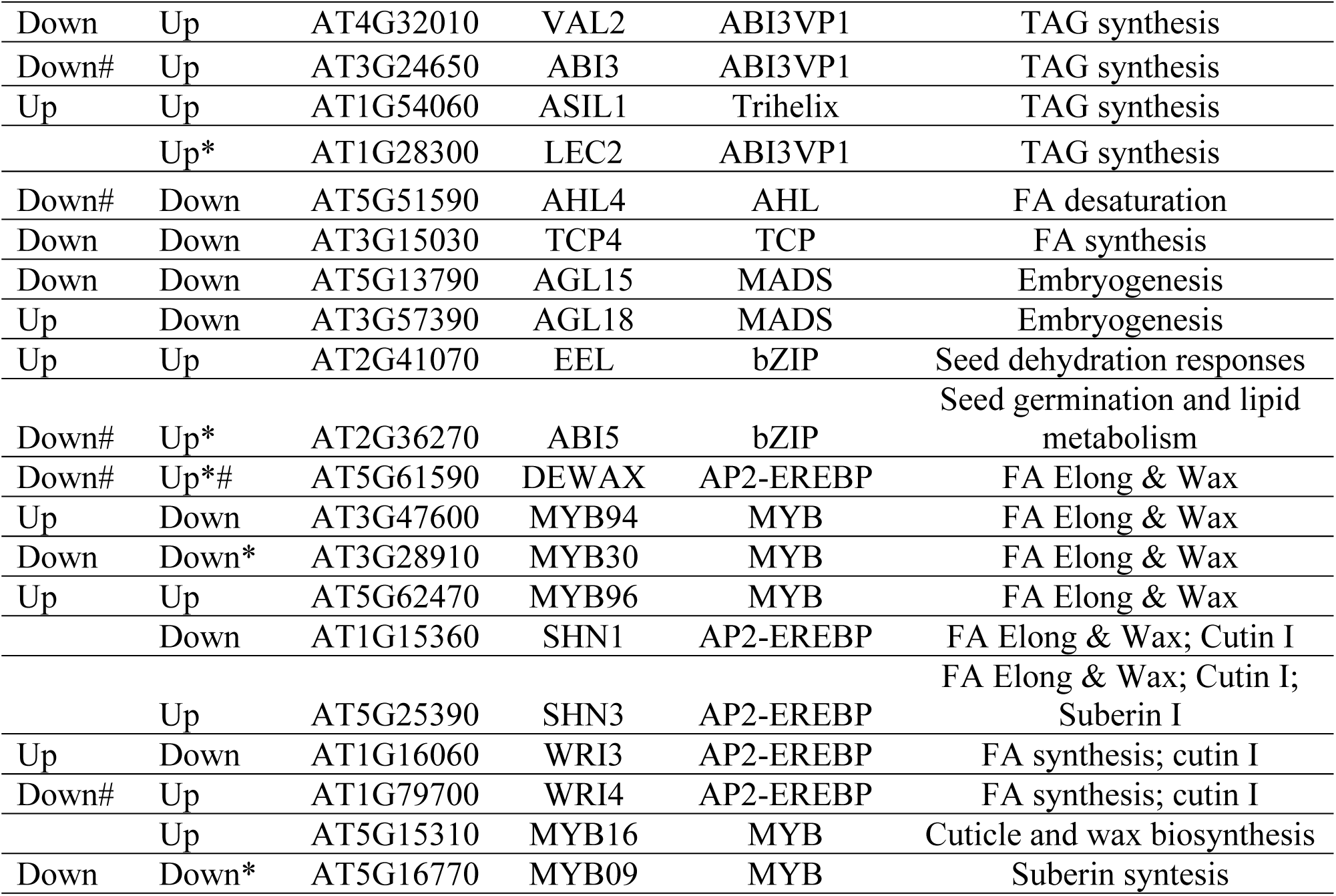
Transcription factors differentially expressed in *badc1/3* lines. *q-value ≤ 0.05, # log2foldchange ≥1 or ≤ −1.

### Omics data supports increased cytosolic acyl-CoA, storage lipid, and protein in *badc1/3*

Once in the cytoplasm, FAs are esterified to CoA and interact with cytosolic ACBPs. ACBP6 (At1g31812), which play a major role in cytosolic acyl-CoA pool maintenance during seed development (Hsiao et al., 2014), was increased at protein level (Fig. 3). Further supporting increased cytosolic acyl-CoA levels, the ABC transporter ABCA9 (At5g61730) (Fig. 3), which is a membrane-attached transporter involved in FA uptake into the ER and is expressed in developing seeds during TAG accumulation (Kim et al., 2013; Li et al., 2016), was upregulated at the transcript level in the *badc1/3* developing seeds. Although cytosolic acyl-CoA levels were not measured in our analysis, upregulated ACBP6 along with upregulated ABCA9 suggest increased processivity of cytosolic acyl-CoA for lipid metabolism in the ER.

**FIGURE 3.**
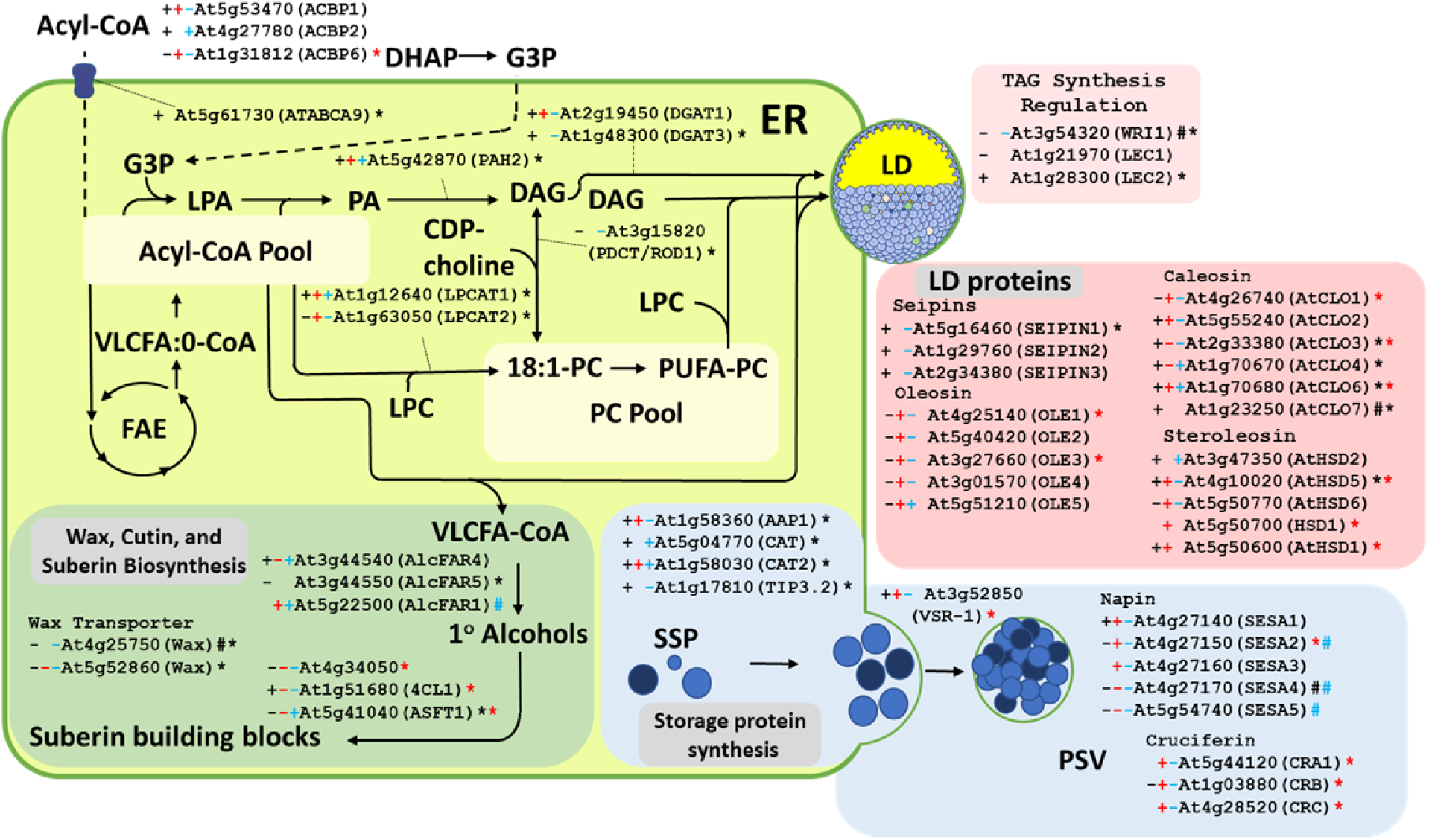
Proteins, transcript, and translation efficiency changes in TAG biosynthesis and seed reserve storage proteins. The plus and minus signs represent upregulation and downregulation respectively in the *badc1/3* line. The black, red, and blue color represent transcript, protein, and translation efficiency data, respectively. The number sign represents Log2FC ≥1 or ≤ −1 for transcript and protein, the asterisk represents genes with q value ≤ 0.05 for transcript and pvalues ≤ 0.05 for protein, these are also color-coded. The blue number sign represents translation efficiency Z-score ≥ 2 or ≤ −2. Abbreviations: acyl-CoA binding protein (ACBP); very long chain fatty acids (VLCFA); fatty acid elongation (FAE); phosphatidate phosphatase (PAH2); 1-acylglycerol-3-phosphocholine acyltransferase, lysophospholipid acyltransferase (LPCAT); acyl-COA:diacylglycerol acyltransferase (DGAT); phosphatidylcholine:diacylglycerol cholinephosphotransferase (PDCT/ROD1); amino acid permease 1 (AAP1); cationic amino acid transporter (CAT); beta-tonoplast intrinsic protein (TIP3.2); wrinkled 1(WRI1); leafy cotyledon (LEC1/2); ATP-binding cassette A9 (ATABCA9); oleosin (OLE); Caleosin (AtCLO); steroleosin (AtHSD); alcohol-forming fatty acyl-CoA reductases (AlcFAR); 4-coumarate:COA ligase (4CL1); aliphatic suberin feruloyl-transferase (ASFT1); wax transporter (Wax); Cruciferin (CB); Napin (SESA)

Lipid assembly steps were also enhanced, including incorporation of FA into PC (a key intermediate in TAG biosynthesis) (Bates, 2022) and TAG (Fig. 3). Lysophosphatidylcholine acyltransferase 1 (LPCAT1, At1g12640), involved in incorporation of newly synthesized FA into PC (Karki et al., 2019), exhibited elevated transcript and protein levels, as well as TE. Phosphatidate phosphatase (PAH2, At5g42870), responsible for the conversion of PA into DAG (Eastmond et al., 2010), was upregulated at transcript level. Regarding ER TAG synthesis, diacylglycerol acyltransferase 3 (DGAT3, At1g48300), a cytosolic DGAT (Ayme et al., 2018), was upregulated at transcript level, and DGAT1 (At2g19450), the primary enzyme involved in TAG biosynthesis (Lu et al., 2003), was upregulated at transcript, and protein levels, though not statistically significant. This points to increased TAG production consistent with [^14^C]acetate pulse/chase data demonstrating increased flux through PC for slightly increased flux of FA into TAG at 9-10 DAG (Fig. 1D, E). These data suggest the rate of TAG synthesis is slightly increased in the *badc1/3* mutant, and most likely the increased oil content in mature seeds is the result of accumulation over time during seed filling.

Increased TAG accumulation was supported by upregulated seipins (At5g16460, At1g29760) at the transcript level, and lipid droplet-associated proteins (LDAPs), oleosin isoforms 1 through 5 (At4g25140, At5g40420, At3g27660, At3g01570, At5g51210), steroleosins (At4g10020, At5g50770, At5g50700, At5g50600) and caleosins (At4g26740, At2g33380, At1g70680) at the protein level. However, the TE of oleosins, caleosins, and steroleosins was decreased. Increased lipid droplet protective proteins may be another metabolic adaptation to prevent lipid turnover and accommodate increased oil content in the *badc1/3* mutant (Kambhampati et al., 2021, 2019). However, reduced TE may imply cellular mechanisms to prevent over-drafting of ribosome complexes for extremely abundant transcripts, or a metabolic adaptation to resist excessive TAG accumulation by limiting LDAPs availability.

The stimulated changes in triacylglycerol accumulation in the mutant had negative compensatory effects on metabolic pathways for wax, cutin and suberin formation. The acyl-activating enzyme, LACS2, reported recently as important for cuticle permeability (Xie et al., 2020), was downregulated at both transcript and protein levels in the mutant developing seeds. Genes involved in wax, cutin, and suberin biosynthesis were also downregulated (Fig. 3, and table 1). This is interpreted as an attempt to shift FA utilization from surface lipids to TAG, and may account for the increased amount of elongated (≥20C) FA in the seed oil (Supplemental Fig. S1).

Consistent with the increase in total seed protein (Fig. 1), seed storage proteins, including napins (At4g27140, At4g27150, At4g27160) and cruciferins (At5g44120, At1g03880, At4g28520), were increased (Fig. 3). Storage protein accumulation is further supported by increases in amino acid transporters amino acid permease 1 (AAP1, At1g58360) and cationic amino acid transporter (CAT, At5g04770) (Fig. 3), although a number of amino acid biosynthetic steps and those pertaining to nitrogen metabolism were decreased, as described in Figure 3 and Supplementary Figure 8.

### Increased lipid mobilization by upregulated lipases, β-oxidation, and the glyoxylate cycle

To investigate whether the increased levels of lipids were a result of alteration in lipid turnover, expression of genes involved in TAG mobilization (Fig. 4), including SDP1 lipase (At5g04040), and core genes involved in β-oxidation were examined. Increases in SDP1, acyl-CoA oxidases (ACXs, At4g16760, At5g65110, and At1g06290), multifunctional proteins (At3g06860, At4g29010), and 3-ketoacyl-CoA thiolase (KAT2, At2g33150) at the transcript, protein, and TE levels suggest that in the face of the greater accumulation of lipids, cells were prepared for enhanced turnover of acyl chains. Other steps in the beta oxidation and the glyoxylate cycle were enhanced, including peroxisomal acyl-activating enzymes (At3g05970, At5g27600, At5g16370 and At5g23050), NAD-malate dehydrogenase (PMDH1, At5g09660), catalases (CAT, At1g20620 and At1g20630), citrate synthase (CSY, At3g58750 and At2g42790), and the two glyoxylate cycle exclusive enzymes isocitrate lyase (ICL, At3g21720) and malate synthase (MLS, At5g03860) (Fig. 5).

**FIGURE 4.**
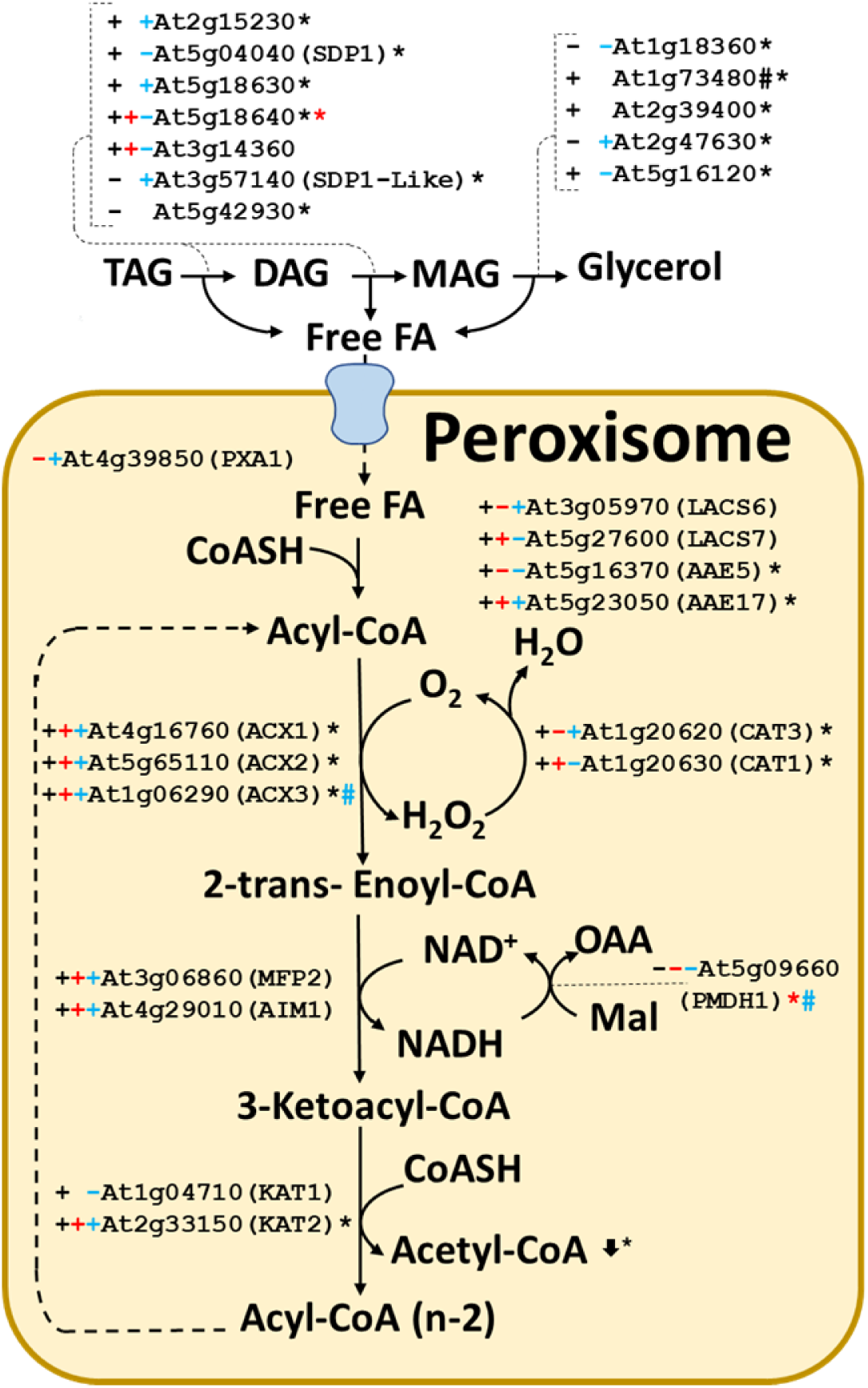
Proteins, transcript, and translation efficiency changes related to TAG mobilization and β-oxidation. The plus and minus signs represent upregulation and downregulation respectively in the *badc1/3* line. The black, red, and blue color represent transcript, protein, and translation efficiency data, respectively. The number sign represents Log2FC ≥1 or ≤ −1 for transcript and protein, the asterisk represents genes with q value ≤ 0.05 for transcript and p or q values ≤ 0.05 for protein while the number sign represents Log2FC ≥1 or ≤ −1 for transcript and protein, these are also color-coded. The blue number sign in blue represents translation efficiency Z-score ≥ 2 or ≤ −2. The number sign in blue represents translation efficiency Z-score ≥2 or ≤ −2. Abbreviations: sugar-dependent1 (SDP1); peroxisomal ABC transporter 1 (PXA1); long-chain acyl-CoA synthetase (LACS); acyl activating enzyme (AAE); acyl-CoA oxidase (ACX); catalase (CAT); multifunctional protein (MFP2/AIM1); 3-ketoacyl-coa thiolase (KAT); peroxisomal NAD-malate dehydrogenase (PMDH1)

### *badc1/3* mutant has altered levels of central metabolites and amino acids

Metabolite pools were measured by LC-MS/MS (Fig. 2 and 5, black up and down arrows next to metabolites) according to Koley et al., (2022). Metabolite measurements demonstrated decreased levels of glycolysis intermediates (glucose 6-phosphate, fructose 6-phosphate, dihydroxyacetone phosphate, glyceraldehyde 3-phosphate and pyruvate), reductive pentose phosphate pathway intermediates (erythrose 4-phosphate, ribulose 1,5-bisphosphate and sedoheptulose 7-phosphate), and TCA intermediates (citrate, iso-citrate, alpha-ketoglutarate, succinate, fumarate, and malate). However, most of the changes were not statistically significant and were consistent with reductions in sucrose, fructose and ADP-glucose levels in the *badc1/3* mutant. Amino acid level changes were similarly marginally altered in the *badc1/3* mutant, including plastid-synthesized amino acids valine, leucine, isoleucine, tryptophane, tyrosine, phenylalanine and histidine. Of these, histidine was statistically significant. The levels of amino acids considered to be synthesized outside of the plastid, including aspartate, asparagine, threonine, alanine, glutamate, glutamine, proline, and arginine, were reduced in the *badc1/3* mutant. In contrast, lysine and methionine were increased. Acetyl-CoA, which is a building block for leucine, was reduced in the *badc1/3* mutant (Fig. 5). Given the increase in total protein (Fig. 1, 3), these results suggest enhanced utilization of amino acids for protein synthesis.

**FIGURE 5.**
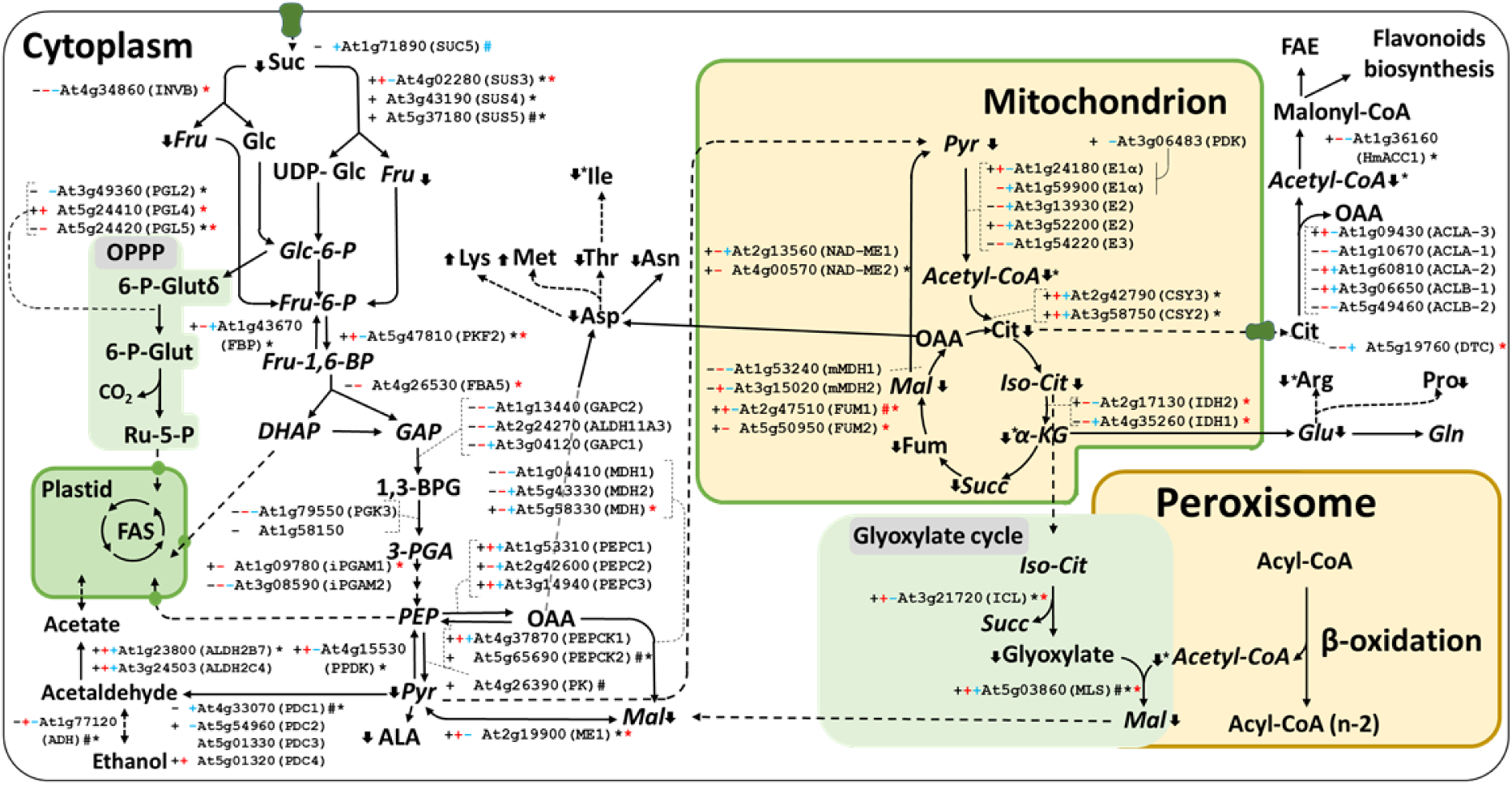
Global proteins, transcript, and translation efficiency changes in cytosolic, mitochondrial, and glyoxylate cycle enzymes caused by *badc 1/3* knockout in developing Arabidopsis seeds. The plus and minus signs represent upregulation and downregulation respectively in the *badc1/3* lines. The black, red, and blue color represent transcript, protein, and translation efficiency data, respectively. The number sign represents Log2FC ≥1 or ≤ −1 for transcript and protein, the asterisk represents genes with q value ≤ 0.05 for transcript and p values ≤ 0.05 for protein, these are also color-coded. The blue number sign represents translation efficiency Z-score ≥ 2 or ≤ −2. The black and green arrows represent decreased and increased metabolite levels, respectively. The asterisk next to the arrow represents p values ≤ 0.05 for the difference in metabolite levels between *badc1/3* mutant and WT. Abbreviations: sucrose-proton symporter 5 (SUC5); sucrose synthase (SUS); invertase (INVB); 6-phosphogluconolactonase (PGL); cytosolic fructose-1,6-bisphosphatase (FBP); phosphofructokinase 2 (PKF2); glyceraldehyde-3-phosphate dehydrogenase (GAPC2); cytosolic-nad-dependent malate dehydrogenase (MDH); phosphoenolpyruvate carboxylase (PEPC); phosphoglycerate kinase (PGK); 2,3-biphosphoglycerate-independent phosphoglycerate mutase (iPGAM); phosphoenolpyruvate carboxykinase (PEPCK); malic enzyme (ME); NAD-dependent malic enzyme (mMDH); fumarase (FUM); citrate synthase (CSY); isocitrate dehydrogenase (IDH2); ATP-citrate lyase (ACLA/B); homomeric acetyl-CoA carboxylase (HmACC); isocitrate lyase (ICL); malate synthase (MSL)

### *badc1/3* mutant has altered carbohydrate transport and organic acid metabolism

In the *badc1/3* mutant seeds, sucrose transporter sucrose-proton symporter 5 (SUC5, At1g71890), a transporter with documented importance for TAG accumulation (Pommerrenig et al., 2013), was upregulated at TE levels. Additionally, sucrose synthases (SUS, At4g02280, At3g43190, and At5g37180), were upregulated (Fig. 5). Increased sucrose transport suggests a heightened demand for glycolysis intermediates in developing seeds in *badc1*/3 mutant, further supported by upregulated cytosolic isoforms of phosphofructokinase (PKF2, At5g47810) at the transcript and protein levels and pyruvate kinase (PK, At4g26390) at the transcript level. Gluconeogenesis fructose-1,6-bisphosphatase (FBP, At1g43670), phosphoenolpyruvate carboxykinase (PEPCK, At4g37870, At5g65690), and pyruvate orthophosphate dikinase (PPDK, At4g15530) were also upregulated at the transcript level. Increased levels for genes involved in cytosolic glycolysis and gluconeogenesis indicated an elevated demand for glycolytic intermediates.

Organic acid metabolism was altered in the *badc1/3* mutant (Fig. 5). Citrate synthase (CSYs At2g42790 and At3g58750) was upregulated at the transcript, protein and TE levels. Citrate can be exported to the cytoplasm and cleaved by citrate lyase generating cytoplasmic acetyl-CoA (Fatland et al., 2002; 2005). In the cytoplasm, acetyl-CoA can be used by HmACCase (At1g36160), which was upregulated at the transcript level, and incorporated into fatty acid elongation or flavonoid biosynthesis. Indeed, the *badc1/3* mutant demonstrated increased fatty acid elongation (Supplementary Fig. S1), while omics data suggested flavonoid biosynthesis is mostly downregulated (Supplementary Fig. S9), pointing to yet another metabolic response to increased TAG content in *badc1/3* mutant. Glyoxylate cycle was increased in *badc1/3* in isocitrate lyase (ICL, At3g21720), and malate synthase (MLS, At5g03860) (Fig. 5). Malate can be converted into pyruvate by malic enzyme (ME) in the cytoplasm (At2g19900) and mitochondria (At4g00570), both upregulated in the transcript and protein datasets. Metabolite measurements of malate and pyruvate pointed to reduced levels of said organic acids, demonstrating a possible pull for pyruvate (Fig. 5). We hypothesized that in *badc1/3* mutant, the glyoxylate cycle pyruvate is likely pulled into storage proteins (Fig. 1B) through pyruvate derived AA, and is reincorporated into FA through pyruvate conversion into acetate, then plastidial acetyl-CoA which could be used by ACCase for FA synthesis and accumulation as TAG (Fig. 6).

**FIGURE 6.**
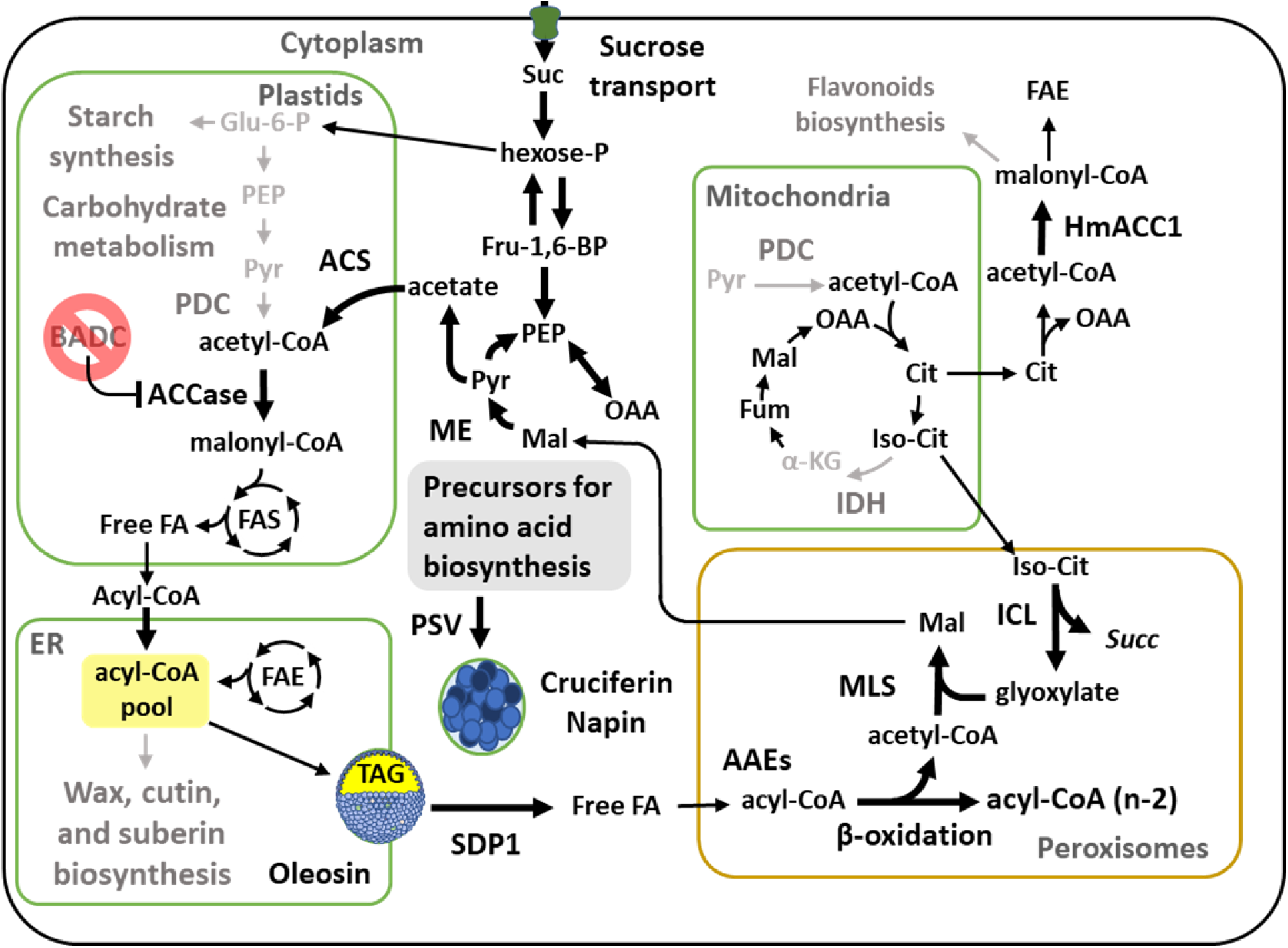
Summarized description of metabolic change in *badc1/3* mutant developing seeds. Knocking out BADC1 and BADC3 results in increased FAS, which is counteracted by reduction of acetyl-CoA provision from PDC. The resulting reduction in plastidial acetyl-CoA can be compensated by ACS. Plastid synthesized FAs are exported into the cytoplasm and into the ER by upregulated ABCA9. In the ER, acyl-CoA accumulation into extracellular surface lipids is reduced, and more acyl-CoA can be incorporated into TAG. The increased TAG content requires increased oleosin abundance, which is responded to by restricting oleosin availability. As another response to limit increased oil, TAG conversion into organic acid is increased by upregulated SDP1, β-oxidation and glyoxylate cycle. Citrate can provide cytosolic acetyl-CoA to be used by HmACCase and reincorporated into lipids through fatty acid elongation (Supplementary Figure 1). While malate can be converted into pyruvate, which can be converted into acetate and reincorporated into acetyl-CoA by ACS. Therefore, resulting in the reincorporation of carbon into TAG. Grey color indicate downregulation, while larger black arrows indicate upregulation.

## DISCUSSION

The hetACCase is a promising target for metabolic engineering to improve seed oil content due to its key role in *de novo* FAS (Chapman & Ohlrogge, 2012). However, efforts to increase hetACCase activity have only led to incremental increases in seed TAG content (Salie et al., 2016; Ye et al., 2020; Wang et al., 2022). This could be due to unexpected mechanisms regulating the balance between *de novo* FAS and other pathways that share acetyl-CoA as either a precursor or product. Understanding this metabolic crosstalk could not only explain the limitations of increasing seed oil, but also identify potential targets for biotechnology enhancement of seed oil. Previous parallel omics studies that investigated higher seed oil content were applied towards near-isogenic lines, which contain additional background gene lesions that may have contributed to the observed transcriptome and proteome alterations (Hajduch et al., 2010; Lin et al., 2006). In this study, the molecular and biochemical consequences were thoroughly studied employing a defined mutant (*badc1/3*) with deregulated hetACCase activity that resulted in enhanced seed oil accumulation (Fig. 1).

### The *badc1/3* mutation increases seed oil and protein with limited increases in the rate of FAS

Consistent with previous studies (Keereetaweep et al., 2018), we confirmed that the *badc1/3* mutant line has increased seed oil content (Fig. 1A). We also documented the surprising increase in total seed protein (Fig. 1B, C). Given the well-established inverse relationship between oil and protein within plant seed (Kambhampati et al., 2019; Wilcox and Shibles, 2001; Wilcox, 1998; Hymowitz et al., 1972; Johnson et al., 1955), this finding was unexpected. Our multi-omics data suggest that increasing hetACCase activity in the *badc1/3* mutant leads to changes in gene expression and cognate enzymes involved in carbohydrate, organic acid, fatty acid, and acetyl-CoA metabolism, resulting in increased oil and protein content in mature seeds. Of note, starch biosynthesis, wax biosynthesis, and flavonoid biosynthesis were each downregulated in *badc1/3*. Although these compounds were not measured in our analysis, we speculate carbon is being shifted from the synthesis of starch, extracellular lipids, and flavonoids towards storage oil and protein. Most curiously, [^14^C]acetate labeling, in developing seeds, suggested only a slight increase in FAS. Nevertheless, the ∼17% increase in seed TAG accumulation (Fig. 1A), could be explained by the small increase in FAS if sustained over the entire two-week period of TAG accumulation in Arabidopsis seeds.

### *badc1/3* mutation may increase sink capacity and alter glycerolipid metabolism

In addition to the slight increase in FAS, comparative omics analyses revealed changes consistent with increased sink capacity and carbon partitioning towards storage lipids in *badc1/3* developing seed. Upregulation of the plasma membrane sucrose transporter SUC5 suggested enhanced sucrose uptake by seeds (Fig. 5). Increased expression of the fatty acid transporter ABCA9 is consistent with enhanced FA processivity in the ER (Fig. 3) (Kim et al., 2013; Pommerrenig et al., 2013). Overexpression of ABCA9 resulted in a 40% increase in oil content in seeds but did not change overall protein and carbohydrate levels in Arabidopsis, indicating that ABCA9 is most likely involved in partitioning towards storage lipids by augmenting FA transport into the ER (Kim et al., 2013). Upregulation of these two transporters, which are known or suspected to be involved in TAG accumulation, is another metabolic adaptation to accommodate increased *de novo* FAS in *badc1/3*.

In the ER, FAs have multiple fates, including (1) membrane lipid assembly, (2) acyl exchange in/out of PC for further modifications, and (3) incorporation into TAG for lipid storage with PC-derived DAG as the predominant precursor for TAG synthesis (Bates, 2016). The initial increase in [^14^C]PC labeling in the *badc1/3* line, followed by enhanced TAG labeling during the chase (Fig. 1D, E) is consistent with enhanced acyl flux through PC prior to TAG biosynthesis in the mutant. Additionally, it indicates that despite the 17% increase in TAG, there was no major change to the acyl assimilatory pathway for TAG biosynthesis in the *badc1/3* mutant. Two changes revealed by comparative omics are consistent with enhanced flux of acyl groups through PC prior to TAG biosynthesis. Phosphatidate phosphohydrolases are responsible for the conversion of PA into DAG and may regulate phospholipid and TAG synthesis (Eastmond et al., 2010). Therefore, upregulated phosphatidate phosphatase (PAH2) may indicate enhanced DAG production for PC and subsequent PC-derived DAG and TAG biosynthesis (Fig. 3). Lysophospholipid acyltransferase (LPCAT1), which is involved in incorporation of nascent acyl-CoA into PC through acyl editing (Karki et al., 2019) is also upregulated at the transcript and protein levels. Upregulation of FA synthesis has been demonstrated to increase rates of PC acyl editing (Zhou et al., 2020). Regarding ER TAG synthesis, upregulated DGAT1, though not statistically significant, points to some increase in TAG production, and is consistent with [^14^C]acetate pulse/chase data demonstrating slightly increased flux of FA into TAG (Fig. 1D, E).

### Increased TAG accumulated in *badc1/3* mutant is likely counteracted by arrested lipid droplet development and enhanced lipid turnover

Our multi-omics data revealed oleosins and caleosins were both up- and down-regulated at protein and TE levels, respectively (Fig. 3). This suggested limited transcription of oleosins and caleosins could be another adaptation to counterbalance increased storage lipid accumulation in *badc1/3*. This aligns with the role of LDAPs (both oleosins and caleosins) in stabilizing lipid droplets (Shao et al., 2019; Winichayakul et al., 2013), therefore, accommodating TAG accumulation. Reduced TE of oleosins and caleosins suggests a metabolic adaptation to resist excessive TAG accumulation by limiting LDAP availability, which is further supported by increased lipid mobilization in the *badc1/3* mutant.

Storage lipid mobilization involves the breakdown of lipid droplets by lipases, resulting in the release of free FAs that are imported into peroxisomes and further catabolized by β-oxidation. SDP1 is the major lipase demonstrated to be fundamental for TAG hydrolysis in many species (Azeez et al., 2022; Aznar-Moreno et al., 2022; Kelly et al., 2011; O’Hara et al., 2006). Upregulation of SDP1 in *badc1/3* seeds showed another layer of controlled regulation resisting excessive TAG accumulation (Fig 4). Additionally, the enzymes involved in β-oxidation, ACX1-3 (Adham et al., 2005; Eastmond et al., 2000a;; Pinfield-Wells et al., 2005) and KAT2 (Germain et al., 2001) were upregulated at the transcript, protein and TE levels. This suggests β-oxidation is enhanced in the *badc1/3* mutant. Lastly, upregulated glyoxylate cycle enzymes ICL (Eastmond et al., 2000b) and MLS (Cornah et al., 2004) reflect the increased need for metabolizing peroxisomal acetyl-CoA, and further corroborates the enhancement of β-oxidation in the *badc1/3* mutant. This is consistent with a hypothesized feedback mechanism that has been suggested to occur between peroxisome and lipid droplets via SDP1 activity during seedling establishment to control the optimal supply of free FAs to the peroxisomal β-oxidation process (Kataya et al., 2019). Despite the increase in FA β-oxidation, largescale TAG turnover as not detected (Fig. 1), and thus the indicated TAG/FA turnover limits the total TAG accumulation rather than decreasing total TAG levels. Future crop engineering efforts that utilize deregulation of ACCase may also need to control acyl turnover to further enhance seed oil.

### Deregulating ACCase leads to metabolic adjustments to counterbalance the increased lipid synthesis

Upregulation of sucrose synthase, and genes involved in cytosolic glycolysis and gluconeogenesis suggest an increased demand for glycolytic intermediates such as pyruvate, triose- and hexose-phosphate. However, metabolomics data showed lower levels of these metabolites (Fig. 2, 5), indicating that in *badc1/3* seeds, glycolysis intermediates were likely pulled to be used as precursors with metabolic pathways at a higher rate, and, therefore, are at lower levels in the *badc1/3* mutant. The *badc1/3* mutant showed upregulation of cytosolic glycolysis/gluconeogenesis in contrast to downregulated plastid glycolysis, which could reflect a partial switch in metabolism from storing carbon in lipid-form towards the production of organic acids (TCA intermediates), which can be employed as carbon skeletons for amino acid production and storage protein accumulation during seed filling. Thus, changes in gene expression and enzyme levels may reflect a metabolic adaptation to limit the increased oil caused by *badc1/3* mutation and redirect some carbon to precursors for the synthesis of amino acids. Surprisingly, our omics data suggested reduction of amino acid biosynthesis (Supplementary Fig. S8), thus contrasting carbon being allocated towards storage protein. Nevertheless, increased storage protein content in both developing (Fig. 3) and mature seed (Fig. 1) clearly demonstrates there is enhanced carbon and nitrogen accumulation in the form of storage protein. Thus, the observed multi-omic changes in amino acid biosynthetic pathways could also be a regulatory adaptation in attempt to slow down the enhanced protein biosynthesis occurring in the *badc1/3* mutant.

Plastid glycolysis and the plastid pyruvate dehydrogenase complex (pPDC) are the main source of acetyl-CoA for FAS (Mooney et al., 1999; Schwender et al., 2004). Transcriptional regulation appears to limit the supply of acetyl-CoA feeding hetACCase activity, as indicated by the broad range transcriptional downregulation of plastid translocators (glucose 6-phosphate/phosphate transporter 2, triose-phosphate∕phosphate translocator, and phosphate/PEP translocator) and plastidial glycolytic enzymes (phosphoglucose isomerase 1, phosphofructokinase 5, fructose-bisphosphate aldolase, glyceraldehyde 3-phosphate dehydrogenase, and phosphoglycerate kinase), as well as each subunit to the pPDC.

De-regulation of hetACCase activity, together with reduction in the pPDC would lead to depletion of plastid acetyl-CoA, which contradicts higher seed oil observed in *badc1/3.* However, plastid acetyl-CoA can also be supported by plastid ACS (which was highly upregulated in *badc1/3*) since acetate is a viable source of carbon for *de novo* FAS (Lin and Oliver, 2008), The contribution of ACS and pPDC for *de novo* FAS has been researched extensively (Liedvogel and Stumpf, 1982; Murphy and Stumpf, 1981; Bao et al., 2000), although never in a high-oil mutant. In our data, ^14^C labeling from [^14^C]acetate, showed increased incorporation of ^14^C into FAs in *badc1/3* seedlings (Supplementary Fig. S2). In mature plants, acetate is thought to originate from the oxidation of ethanol and the non-oxidative decarboxylation of pyruvate (Fu et al., 2020). Both sources involve the formation of acetaldehyde, which is converted into acetate by aldehyde dehydrogenases (Wei et al., 2009), and can provide an alternative route for carbon incorporation into FA, that bypasses pPDC (Mellema et al., 2002). Our data showed two upregulated aldehyde dehydrogenases, indicating a putative increase in acetate production and corroborating a possible partial shift of carbon source for FAS from pyruvate to acetate. This shift, over the course of seed development, can contribute to FAS and subsequent oil accumulation in the *badc1/3* mutant. Considering that acetate availability can influence ACS contribution to FAS, as demonstrated in *C. reinardtii* (Fan et al., 2012; Goodson et al., 2011; Ramanan et al., 2013), ACS may be a potential target for enhanced FA synthesis in both WT and BADC mutants. In the *badc1/3* mutant, [^14^C]acetate pulse-chase in developing seeds showed a slightly increased rate of TAG accumulation and comparative omics supported increased TAG accumulation (increased LDAPs). Conversely, enhanced lipid mobilization was also observed (upregulated SDP1, β-oxidation, and glyoxylate cycle). We conclude the *badc1/3* mutation resulted in both increased FAS and TAG accumulation, the latter being limited by a futile cycle involving metabolic responses to apparently limit excessive oil accumulation (Fig 6). The increase in total seed protein was surprising but supported through multiple independent experiments (Fig. 1B, C, and Fig. 3). Together these results indicate that metabolic adaptations as a result of the de-regulation of hetACCase are complex and that future engineering of very high oil seed will likely need to address not only the activity of ACCase but also metabolic adaptations that limit enhanced TAG accumulation.

## EXPERIMENTAL PROCEDURES

### Plant material and growth conditions

Arabidopsis thaliana Col-0 and *badc1/3* T-DNA insertion mutant crosses were grown in a growth chamber under 16 h day length with 100-150 µmol photons m-2 sec-1 white light measured at the soil surface. Temperatures ranged from 21-23°C during the day and 18-20°C at night. Individual plants (seed storage compounds) or four plants (expression, proteomics, and labeling) were grown to maturity in 8 cm x 8 cm pots in SunGro potting mix LC1 (70% peat, 20% pumice, and 10% sand; SunGro Horticulture, Agawam, MA, USA). Plants were fertilized weekly with 500 ppm Peters 20:15:20 (J.R. Peters) until harvest. Siliques were collected at 9-10 days after flowering (DAF) (Ruuska et al., 2002).

### Seed protein quantification

Total soluble protein was extracted and quantified from 5 mg Arabidopsis seeds following the protocol described in Wang *et al*., (2022). Seed storage proteins were visualized from 100 µg of seed material on a 12% SDS-PAGE compared to a Precision Plus Dual Color Protein Standard (Bio-Rad, Hercules, CA, USA). Gels were stained for 1 hr using a 0.1% Coomassie Brilliant Blue, 40% methanol, and 10% glacial acetic acid solution (Sigma Aldrich) and an orbital shaker (60 rpm) (Wang et al., 2022). Destaining was completed with successive washes of 40% methanol and 10% glacial acetic acid solution while shaking, and gels were imaged with a Syngene G-Box imaging system (Chemi-XT4; Syngene, Bangalore, India).

### Lipid analysis

Seed quantification was performed by first counting and weighing 100 seeds which were then analyzed according to a modified method based on Li *et al*. (2006). First, samples were combined with 1.0 mL freshly prepared 5% (v/v) sulfuric acid in MeOH and 200 µL of toluene with 0.0005% (w/v) butylated hydroxy toluene (BHT; Sigma Aldritch) as well as 40 mg tripentadecanoin (15:0; Nu-chek Prep, Elysian, MN, USA) as an internal standard in a 10 cm x 1 cm glass tube. Tubes were sealed with PTFE-lined caps and incubated at 85°C for 1.5 h. Samples were cooled and 1 mL of hexane and 0.8 mL 0.88% (w/v) potassium chloride (KCl; Sigma Aldrich) was added. Samples were then vortexed and centrifuged at 3,000 × *g* for 5 min to achieve phase separation. FAME extracts were analyzed from the upper hexane phase with an Agilent model 7890 GC with flame ionization detector (FID) and a DB FATWAX UI column (30 m X 0.25 µm; Agilent). The GC-FID was set to a split ratio of (1:20), with injector and FID temperatures set to 255 °C and helium flow of 1.6 mL/min. The oven temperature program started at 120 °C and ramped to 200 °C at 25 °C/min. Next, the oven was ramped at 10 °C/min to 240 °C and held at that temperature for 3.5 min.

### Arabidopsis pulse/chase seed labeling

Developing Arabidopsis seeds collected at 9-10 DAF were labeled following the protocol described in Kotapati and Bates (2021), with the following changes. Following developing seed collection, 4 replicates of both WT and *badc1/3* were aliquoted into sterile disposable 16×100 culture tubes containing fresh growth media containing (5 mM MES pH 5.8, 0.5% sucrose, 0.5x MS salts, 0.5 mM acetate). Samples were placed into a shaking water bath set to 23 °C under light (100 µE) and equilibrated for 1 hr. Next, growth media was removed, and developing seeds were labeled for 1 hour in 200 µL growth labeling media containing 5 mM MES pH 5.8, 0.5% sucrose, 0.5x MS salts, 0.5 mM [14C] acetate (54.3 mCi/mmol; American Radiolabeled Chemicals Inc, St. Louis, Mo). Following labeling, developing Arabidopsis seeds were washed 3 times with cold growth media and finally re-suspended in 1.25 mL of growth media. Samples were collected at the end of the wash step as well as 1, 5, 16, 24, and 48 h after labeling by pipetting 200 µL of developing seeds and media into pre-weighed (see lipid analysis below) 8 mL glass vial and PTFE-lined cap containing 2 mL hot isopropanol (85 °C). Samples were incubated for 15 min to inhibit lipase activity then allowed to cool to room temperature and stored at −20 °C. After sample collection, the remaining volume of growth media was exchanged to promote continued seed development. Seedlings were kept on a 16 h photoperiod identical to the parent plants and seed labeling was timed to correspond to the middle of that photoperiod.

### Analysis of label accumulation in developing Arabidopsis seeds

Following labeling, developing seeds were first homogenized, then lipids were extracted by adding 6 mL of a 2:1 mixture of chloroform: methanol and vortexing thoroughly. Cell debris was pelleted by centrifuging for 5 min at 2500g and the supernatant was transferred to a new 13 mL glass tube with PTFE-lined cap. The extraction was then repeated, and the supernatants were combined. Next, the remaining tissue was dried in a fume hood, weighed, digested in 0.3 mL solusol (National Diagnostics), and counted in 10 mL of Soluscint (liquid scintillation cocktail; National diagnostics). In parallel, the lipids were extracted from the organic phase by adding 2 mL of 0.88% KCl (Sigma-Aldritch) to force phase separation, vortexing, and centrifuging at 2500x g for 5 min. The lower organic phase was transferred to a new 8 mL glass tube and lipids were back extracted from the upper aqueous phase by adding 2 mL chloroform and repeating centrifugation and transfer steps. Finally, the combined lipid samples were blown down under the N stream and re-eluted in 0.5 mL toluene for analysis. Total radioactivity was measured from both lipid and soluble fractions by aliquoting 10% of each and mixing with 5 and 10 mL of ecoscint A (National diagnostics), respectively. All liquid scintillation samples were counted using a liquid scintillation counter following standard procedures (Tri-Carb 4910TR; Perkin Elmer, Waltham, MA). Radio analysis of ^14^C-acetate labeled lipid species was completed using an Agilent 1260 Infinity HPLC system coupled to a Lab Logic β-Ram6 for in-line radioactivity detection using modifications of the standardized method (Kotapati & Bates, 2020) as described in Wang *et al*., (2022).

### Total RNA isolation for RNA-seq analysis

Total RNA extraction was performed in three biological replicates for each sample using the NucleoSpin® RNA Plant kit (MACHEREY-NAGEL), as recommended by the manufacturer. In short, powdered seeds (9-10 DAF) ground in liquid N2 were added to 350 μL of buffer RAP (1% ß-ME) and vortex vigorously. The lysate was filtered by centrifugation with NucleoSpin® Filter for 1 min at 11,000 x g. The filtered lysate was added 70% ethanol, homogenized, and bonded to NucleoSpin® RNA Plant Column by centrifuging for 30 s at 11,000 x g. Column containing RNA was added of 350 μL MDB (Membrane Desalting Buffer) and centrifugated at 11,000 x g for 1 min to dry the membrane. DNase reaction mixture was prepared according to the manufacturer’s instruction and 95 μL DNase reaction mixture was directly added onto the center of the silica membrane of the column and incubated at room temperature for 15 min. A column with Dnase digested RNA was washed by adding 200 μL Buffer RAW2, 600 μL Buffer RA3, and 250 μL Buffer RA3 in three consecutive centrifugations for 30 s at 11,000 x g. Total RNA was eluded in 60 μL RNase-free H_2_O by centrifugation at 11,000 x g for 1 min. Isolated RNA was stored at −80C and sent to Genomics Technology Core - University of Missouri for RNA sequencing.

### Translating RNA affinity purification RNA sequencing (TRAP-Seq) analysis

Plants used for TRAP-Seq experiments express a transgene consisting of the 6X histidine-FLAG-epitope tagged-ribosomal protein L18 (HF-RPL18) under the control of the glycinin promoter from soybean (Glycine max). HF-RPL18 was cloned into the pBinGlyRed3 vector (provided as a gift from the Dr. Edgar Cahoon lab - University of Nebraska, Lincoln) using the XbaI restriction site. The binary vector, pBinGlyRed3 containing HF-RPL18 was then transformed into Agrobacterium tumefaciens (C58C1) and then transformed into WT (Col-0) and *badc1/3* plants using the floral dip method (Clough and Bent, 1998). Seeds harvested from the resulting plants (T1) were screened for DsRed fluorescence and expression of RPL18 protein using commercial Anti-FLAG antibody (MilliporeSigma, Burlington, MA, USA).

Enrichment of the ribosome-associated RNA was according to previously described methods (Castro-Guerrero et al., 2016; Kimberlin et al., 2021; Zanetti et al., 2005). Whole siliques (9-10 DAF) were used as starting tissue materials but the specificity of the glycinin promoter-driven HF-RPL18 expression in seeds allowed the exclusion of transcripts from the siliques. Flash-frozen tissue was ground using a mortar and pestle in liquid N_2_ yielding approximately 10–15 g of powdered tissue per sample. To this, chilled ribosome extraction buffer (200 mM Tris-HCl pH 9.0, 200 mM KCl, 25 mM EGTA, 36 mM MgCl_2_, 1 mM DTT, 50 µg mL^−1^ cycloheximide, 50 µg mL^−1^ chloramphenicol, 1 mM PMSF, 1 % igepal CA-630, 1 % Brij 35, 1 % triton X-100, 1 % tween 20, 1 % tridecyl ether, 1 % sodium deoxycholate, 0.5 mg mL^−1^ Heparin) was added at 2:1 ratio (buffer: tissue), and the mixture was rocked gently at 4 °C for 30 min followed by centrifugation at 16,000 x g for 15 min at 4 °C. The supernatant was filtered through a layer of miracloth followed by incubating with 300 µL of EZ view Red Anti-FLAG M2 Affinity Gel (MilliporeSigma, Burlington, MA, USA) for 2 h at 4 °C with gentle rocking. Beads were recovered by brief centrifugation and were washed four times with 10 mL of wash buffer (200 mM Tris-HCl pH 9.0, 200 mM KCl, 25 mM EGTA, 36 mM MgCl_2_, 1 mM DTT, 50 µg mL^−1^ cycloheximide, 50 μg mL^−1^ chloramphenicol). RNA was then extracted from the washed beads using the Direct-zol RNA Miniprep Plus Kit (Zymo Research, CA, USA). Total RNA extraction was performed as described by Nguyen et. al., 2013. Approximately 3 µg each of total and the TRAP RNA samples (100 ng mL^−^1) in three biological replicates each was sent to Novogene Corporation Inc. (Sacramento, CA) for RNA-Seq analysis.

### RNA-seq and TRAP-seq data analysis

In sequencing data analysis, the raw Illumina sequencing data was passed through the protocol of Cufflinks suite tools in Linux (Trapnell et al., 2012). The raw paired reads were mapped to the reference genome (TAIR10) by using Tophat (Trapnell et al., 2009). Gene expression was compared by Cuffdiff (Trapnell et al., 2012). Log2 fold change of FPKM for all identified genes was calculated, with p-values and q-values. The TE was calculated by dividing the TRAP-Seq reads by the RNA-Seq reads (Table S2). The TE change value is calculated by taking the ratio of the TE of *badc1/3* and TE of WT. Z-score normalization was then performed on the log2 transformation of the TE changes, and |Z-score| ≥1 or 2 was included in the figures.

### Protein extraction, quantification, and digestion Sample Processing

Three biological replicates for each developing seed sample were ground in liquid nitrogen and extracted using phenol according to Mooney et. al., (2004). Protein pellets were resuspended in urea buffer (6 M urea, 2 M thiourea, 100 mM ammonium bicarbonate, pH 8.0) and quantified using EZQ (Invitrogen/Life Technologies).

A 1:10 dilution of the urea solubilized protein was made into urea buffer and the protein was quantified by EZQ assay according to the manufacturer’s protocol (Invitrogen/Life Technologies). The assay paper was scanned using the FLA-5000 Fujifilm imager and quantification results were obtained using the Multi Gauge v2.3 software. Equal amounts (25 µg) of each sample were transferred to fresh tubes and all volumes were normalized to 50 µL with urea buffer prior to digestion with trypsin (1 µg) overnight at 37 °C. Peptides purified by C18 tips according to the manufacturer’s protocol (Pierce/ThermoFisher). Purified peptides were then lyophilized and resuspended in 100 mM HEPES-KOH, pH 8.5.

### Peptide TMT10 labeling and high pH fractioning of labeled peptides

Samples were labeled with TMT10plex reagent according to the manufacturer’s (ThermoScientific) instructions with the following modifications. Peptides were resuspended in 100 mM HEPES-KOH, pH 8.5, and half of the total reagent volume (20.5 µL) was used to label the 25 µg digested protein. Following labeling, all samples were combined and lyophilized. Samples were resuspended in 600 µL of 0.1% TFA and then split into two aliquots such that two high pH fractionation columns (Pierce/ThermoFisher, Pistcataway, NY, USA) were loaded at maximum capacity (100 µg protein). Fractionation was conducted according to the manufacturer’s TMT protocol leading to eight fractions plus a flow-through (unbound fraction) in duplicate. All fractions were lyophilized and resuspended in 10 µL of 5% ACN, and 0.1% FA and transferred to autosampler vials.

### Proteomics-based mass spectrometry

Peptides were analyzed by mass spectrometry as follows: a 2 µL injection was made directly onto a 20 cm long x 75 µm inner diameter pulled-needle analytical column packed with Waters BEH-C18, 1.7um reversed phase resin. Peptides were separated and eluted from the analytical column with a gradient of acetonitrile at 300nL/min. The Bruker (Milford, MA, USA) nanoElute system was attached to a Bruker timsTOF-PRO mass spectrometer via a Bruker CaptiveSpray source. LC gradient conditions: Initial conditions were 2%B (A: 0.1% formic acid in the water, B: 99.9% acetonitrile, 0.1% formic acid), followed by 3 min ramp to 17%B. Then 17-25%B over 17 min, 25-37%B over 15 min, 37-80%B over 10 min, hold at 80%B for 10 min, ramp back (2min) and hold at (2min) initial conditions. The total run time was 60 min. MS data were collected in positive-ion data-dependent PASEF mode (Meier et al., 2015) over an m/z range of 100 to 1700. The instrument was calibrated for m/z and ion-mobility prior to acquisition using ESI-L tuning mix (Agilent Technologies, Santa Clara, CA, USA). PASEF and TIMS were set to “on”. One MS and ten PASEF frames were acquired per cycle of 2.4 (∼1MS and 120 MS/MS). The target intensity for MS was set at 20,000 counts/sec with a minimum threshold of 2500 counts/s. The intensity repetition table (default values) was set to On. A charge-state-based rolling collision energy table was used from 76-123% of 42.0 eV. An active exclusion/reconsider precursor method with release after 0.4min was used. If the precursor (within mass width error of 0.015 m/z) was >4X signal intensity in subsequent scans, a second MSMS spectrum was collected. Isolation width was set to 2 m/z (<700m/z) or 3 (800-1500 m/z). The time stepping mode (default values) was used to generate dual peptide fragments for sequence and reporter ion in the same spectrum.

### Database searches (protein identification)

The acquired data were submitted to the PEAKS (Bioinformatics Solutions Inc, Ontario, Canada) search engine for protein identifications. The TAIR11 database was used (48,359 entries; last update 2/2019). Data were searched with trypsin as enzyme, semi-specific, 3 missed cleavages allowed; carbamidomethyl cysteine and TMT10plex as fixed modifications; oxidized methionine and deamidation of N/Q as variable mods; 20 ppm mass tolerance on precursor ions, 0.1 Da on fragment ions. Search results files were first filtered for 0.1% FDR (peptide spectrum-matches false discovery rate) and >1 unique peptide per protein and exported from PEAKS.

### Quantitation

All samples were run using the TMT-10plex (CID/HCD) quantification method in PEAKS, with the following parameters: quantification mass tolerance of 0.1 Da, FDR threshold 1%, and reporter ion type MS2 according to the channels labeled (described above). WT1 served as the reference channel. Data were then filtered as follows: protein FDR ≤1, ≥1 peptide, present in 3 channels, fold-change ≥2.

### Extraction and relative quantitation of central metabolites

Hydrophilic central metabolites were extracted with methanol, chloroform and water as previously described (Ma et al., 2017) with modifications mentioned in (Koley et al., 2022). The aqueous phase extract was run in LC-MS/MS using an AB Sciex triple quadrupole MS system (QTRAP 6500; AB Sciex, Foster City, CA, USA) connected to a Shimadzu HPLC system (UFLC-XR; Shimadzu Corporation, Kyoto, Japan). Amino acids were separated on an Infinity Lab Poroshell 120 Z-HILIC column (2.7 µm, 100 x 2.1 mm; Agilent Technologies, Santa Clara, CA, USA) and detected in the positive ion mode of MS (Czajka et al., 2020; Kambhampati, Li, et al., 2019), while sugar, sugar phosphates and organic acids were separated on an Imtakt Intrada organic acid column (150 x 2 mm, 3 µm; Kyoto, Japan) and detected in negative ion mode (Koley et al., 2022). Relative quantification of the sizes of the metabolite pool between WT and transgenic line was calculated after correction of sample recovery using internal standards (PIPES, norvaline, and ribitol).

## Acknowledgments

We thank John Shanklin for providing the *badc1/3* mutant seeds. This work was supported by National Science Foundation Grant PGRP IOS-1829365.

